# A subset of gut leukocytes have telomerase-dependent “hyper-long” telomeres and require telomerase for function in zebrafish

**DOI:** 10.1101/2022.01.31.478480

**Authors:** Pam S. Ellis, Raquel R. Martins, Emily J. Thompson, Asma Farhat, Stephen A. Renshaw, Catarina M. Henriques

## Abstract

Telomerase, the enzyme capable of elongating telomeres, is usually restricted in human somatic cells, which contributes to progressive telomere shortening with cell-division and ageing. Using the zebrafish model, we show that subsets gut immune cells have telomerase-dependent “hyper-long” telomeres, which we identified as being predominantly macrophages and dendritics (*mpeg1*.*1*^*+*^ and *cd45*^*+*^*mhcII*^*+*^). Notably, *mpeg*^*+*^ macrophages have much longer telomeres in the gut than in their haematopoietic tissue of origin, suggesting that there is a gut-specific modulation of telomerase in these cells.

Moreover, we show that a subset of gut *mpeg*^+^ cells express telomerase (*tert*) in young WT zebrafish, but that the relative proportion of these cells decreases with ageing. Importantly, this is accompanied by telomere shortening and DNA damage responses with ageing and a telomerase-dependent decrease in expression of autophagy and immune activation markers. Finally, these telomerase-dependent molecular alterations are accompanied by impaired phagocytosis of *E*.*coli* and increased gut permeability *in vivo*. Together, our data show that limiting levels of telomerase lead to changes in gut immunity and gut permeability, which, together, are likely contributors to local and systemic tissue degeneration and increased susceptibility to infection with ageing.

**GRAPHICAL ABSTRACT:** 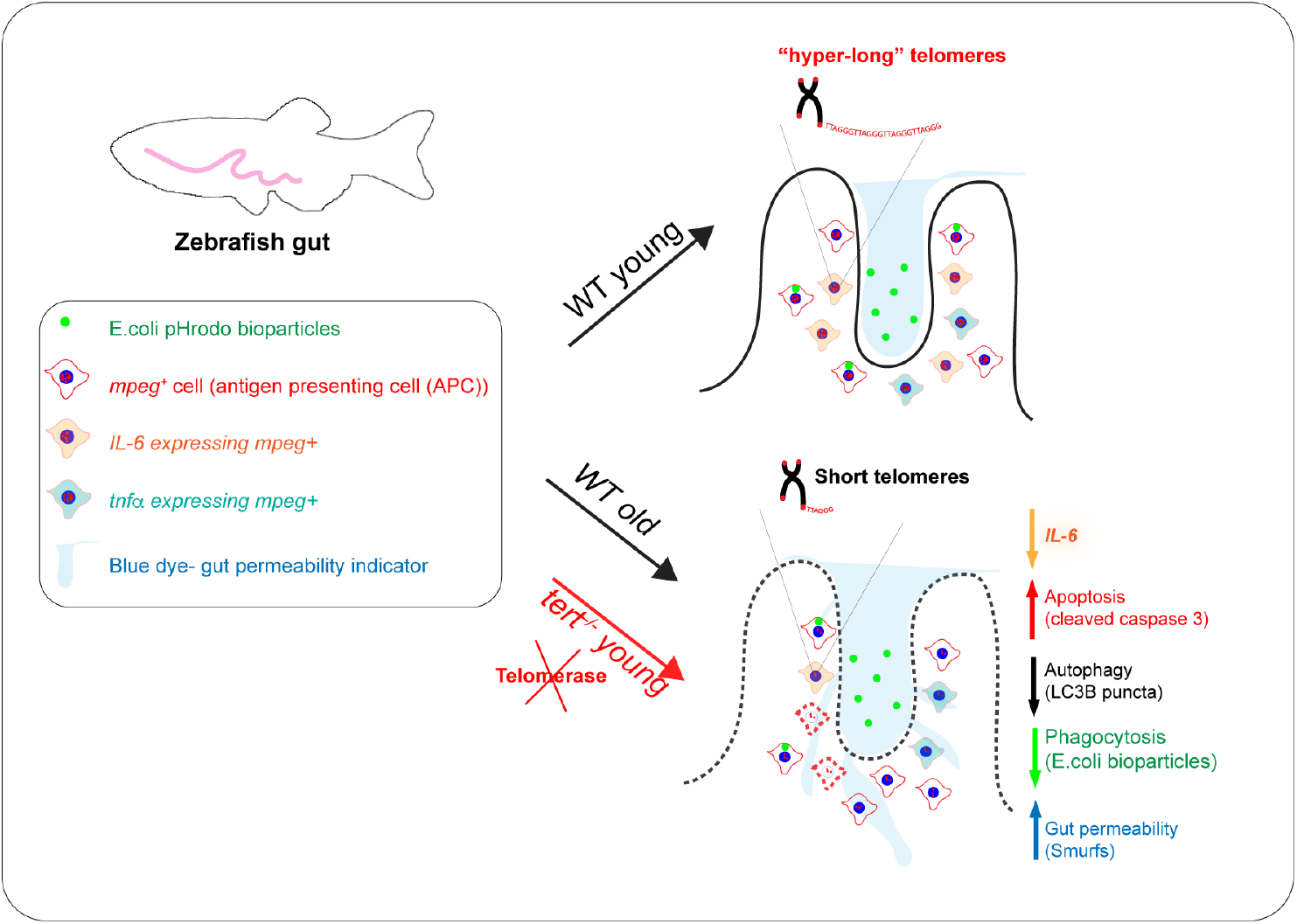

Our data show that limiting levels of telomerase contribute to alterations in gut immunity, namely increased apoptosis, decreased autophagy and immune activation of *mpeg*^*+*^ macrophages in the gut. This is accompanied by a decreased ability to clear pathogens and increased gut permeability. Together, these are likely contributors to local and systemic tissue degeneration and susceptibility to infection with ageing.

## 1 INTRODUCTION

Telomerase, the reverse transcriptase capable of elongating telomeres, is mostly restricted in somatic cells. Telomeres are repetitive DNA sequence of TTAGGG that, together with the protein shelterin complex, provide a protective cap at the end of our chromosomes^1^.

However, in humans, due to limited telomerase expression, and the “end-replication problem”^2^ telomeres shorten with ageing and each cell-division, leading to proliferative exhaustion and replicative senescence^3-7^. This is thought to contribute to the accumulation of cellular senescence with ageing in humans^8^ and senescence has been linked to several age-associated diseases^9^. Telomerase expression is therefore required for the maintenance of germ cells and during the developing embryo, thereby ensuring the replicative potential over generations^10^. Telomerase expression is also a hallmark of tissue stem cells, but its expression in these cells is still insufficient to completely prevent telomere shortening over time^11^. Immune cells are somatic cells that break the rule, as they are capable of modulating telomerase expression in a homeostatic manner. It has long been known that telomerase is up regulated in adaptive immunity, namely in T and B-cells^12^. Telomerase expression in this setting ensures that T-cells undergoing clonal expansion reduce the rate of telomere attrition; thereby ensuring that the replicative potential required for these proliferation outbursts during the adaptive immune response is maintained throughout most of our lives^13-15^. B-cells have also been described to up-regulate telomerase expression during maturation, a process that also relies on proliferation bursts^16^.

Whereas it seems intuitive that a cell type that depends on regular proliferation outbursts for function may have evolved to modulate telomerase expression, such as stem cells, T cells, B cells and most human cancers^17^, it is less obvious why other somatic cells may also do so. Nevertheless this is exactly what a growing body of evidence suggest for a variety of innate immune cells, including neutrophils and macrophages, at least in the context of disease^18-20^. Whether and how telomerase may play a role in innate immune subtypes in the context of natural ageing, in different tissues, however, remains to be determined. Relevantly, telomerase is not only capable of restoring telomeres (canonical function) but also of performing non-canonical functions involved in the regulation of gene expression (recently reviewed here^21^). These so called non-canonical functions of telomerase are important, because they are thought to modulate transcription of genes^22,23^ involved in a wide variety of cellular functions, from DNA repair^24^, to enhanced proliferation^25,26^, inhibition of apoptosis^25^ enhanced migration^18,24,27-31^ and inflammation^32-34^. In the nucleus, these non-canonical functions include transcriptional regulation of genes involved in inflammation, including nuclear factor kappa B (NFKB) and tumour necrosis factor alpha (TNFα)^32-34^, as well as genes involved in cell proliferation^26,35^ and cell survival^36,37^. Telomerase can also translocate to the mitochondria, where it has been shown to play a protective role against DNA damage and oxidative stress^38,39^.

The gut is a key tissue where inflammation must be carefully controlled and modulated to prevent disease^40^. It is the largest immune organ in the body^41^ and is constantly being exposed to foreign antigens, having therefore to maintain a tolerogenic immune status^42^. Perturbations in gut immune regulation are known to contribute to diseases such as inflammatory bowel diseases (IBD) ^43^, ulcerative colitis (UC) and Crohn’s disease (CD)^44^ and indeed ageing-associated degeneration and dysfunction^45-47^, including in zebrafish^48^. Ageing of the gut is an often forgotten ailment in old age that can contribute to malnutrition^49^, ageing-associated anorexia and, consequently, sarcopaenia, frailty, loss of independence and resilience^47,50^. The specific cellular and molecular mechanisms driving changes in inflammation in the gut with ageing, and their relative contribution to the clinical manifestations of an aged gastrointestinal tract, however, is still poorly understood^47^. Previous work in different model organisms, including zebrafish, has highlighted the gut as one of the first tissues to age in a telomerase-dependent manner. Indeed, there is evidence suggesting that the gut plays an important role in the pathogenesis and progression of systemic inflammation, which can lead to multiple organ failure and eventually death^51, 52, 53,54^.

We therefore set out to determine the molecular and functional consequences that of whole organism telomerase depletion may have in gut-associated immune cells in specific. We used the telomerase mutant (*tert*^*-/-*^) zebrafish as a model^55-57^, alongside its WT counterpart. Zebrafish have been previously shown to age in a telomerase-dependent manner, mimicking many aspects of human ageing^55,56,58^. Accordingly, while *tert*^*-/-*^ fish have a lifespan of c. 12-20 months, WT fish typically die between c. 36-42 months of age^55^. Akin to humans, previous work in zebrafish suggested that telomerase expression is likely to be differentially regulated in different cell types, highlighted by the observation that peripheral and kidney marrow blood cells have much longer telomeres than most other somatic cell types in zebrafish^55,56^.

Here, we show that gut immune cells have telomerase-dependent “hyper-long” telomeres, which we identified as being predominantly macrophages/ dendritics (*mpeg1*.*1*^*+*^ and *cd45*^*+*^*mhcII*^*+*^). We show that a subset of gut *mpeg*^+^ macrophages do indeed express telomerase (*tert*) in young WT zebrafish, but that the relative proportion of telomerase-expressing *mpeg*^*+*^ cells decreases with ageing. This is accompanied by telomere shortening and a telomerase-dependent decrease in expression of autophagy (LC3B) and immune activation (IL-6) markers in gut *mpeg*^+^ cells with ageing. Importantly, we show that alongside these telomerase-dependent molecular alterations are accompanied by impaired phagocytosis of *E*.*coli* and increased gut permeability *in vivo*. Together, our data show that limiting levels of telomerase lead to changes in gut immunity and gut permeability likely to contribute to tissue degeneration and susceptibility to infection with ageing^54,59^.

## 2 RESULTS

### 2.1 Gut-associated leukocytes have telomerase-dependent “hyper-long” telomeres, independently of proliferation

Previous work using telomere *in situ* hybridisation (Telo-FISH) in zebrafish gut sections show very distinct cell populations with different telomere lengths, and highlighted a particular subset of cells with “hyper-long” telomeres in the gut, which was absent in the absence of telomerase (*tert*^*-/-*)56^. Together, these observations suggested that, akin to humans, telomerase expression is likely to be differentially regulated in different cell types in WT zebrafish, but the identity of these hyper-long” telomere cells in the gut remained to be determined. Importantly, in terms of telomere length in the gut, the most significant difference between WT and *tert*^*-/-*^ zebrafish at a young age (under 5 months) is the absence of these telomerase-dependent “hyper-long” telomere cells^55,56^. We therefore postulated that these likely constitute a cellular subset particularly dependent on telomerase, and a potential candidate for driving the initial stages of the ageing phenotypes we see developing in *tert*^*-/-*^ and, at later stages, in naturally aged WT. Because it had also been shown that peripheral blood and the head kidney (the bone marrow equivalent in zebrafish), had longer telomeres than other tissues in WT zebrafish^55,56^, we hypothesised that these “hyper-long” telomere cells in the gut were likely to be tissue-associated immune cells. We therefore set out to determine what these cells were, and whether they were still present in the *tert-/-*gut, albeit with shorter telomeres.

Our results confirmed the presence of these “hyper-long” telomere cells in WT young adult zebrafish gut, and that this strong telomere signal (Telo-FISH) was not detected in the absence of telomerase (*tert*^*-/-*^) (**Fig 1A1**). Importantly, we show a clear and complete overlap between these “hyper-long” telomere cells and a pan-immune cell staining (L-plastin) in the gut. Importantly, these gut immune cells are still present in comparable numbers in the absence of telomerase, albeit with much shorter telomeres (**Fig 1A1.1**). To discard the possibility that gut immune cells have longer telomeres due to cell proliferation, which could artificially increase the intensity of the telomeric signal due to DNA replication in S-phase, we performed a control centromere FISH in parallel to the Telo-FISH. For this, we used a fluorescent PNA probe which we designed to be complementary to a near-centromere region that had been shown to be expressed consistently and only once per chromosome (ZEFRFAL1) in the zebrafish genome^60^. If the “hyper-long” telomere cells were replicating, then the “centromeric” signal would also be significantly stronger in these cells. We show that this is not the case (**Fig 1B**). Normalising the telomere signal by the centromeric signal (tel/cent ratio), controls for any differences in DNA content we further normalised our comparisons by normalising the tel/cent ratio of the L-plastin+ cells by the tel/cent ratio of the gut epithelial cells (enterocytes) from the same field of view (FOV). This allowed us to be confident when comparing and pooling the results of multiple FOV from multiple animals together. We detected a clear “hyper-long telomere cell population, which we identified having telomere signal between 20 and 50% stronger than epithelial cells in the gut (1.20 to 1.50 tel/cent ratio in L-plastin^+^ normalised to tel/cent ratio of epithelial cells) (**Fig 1B1**). Again we show that, in the telomerase mutant, all cells have a tel/cent ratio below 1.20. Moreover, as has been shown before^56^, the cell population within the lower telomere intensity, which we identified as epithelial cells based on the well-described nuclear morphology and localisation in the gut villi, has equivalent telomere signal between WT and *tert*^*-/-*^ zebrafish. This again strongly suggests that, as in humans, WT zebrafish have restricted telomerase expression/activity in somatic cells. We further confirm that this increased telomere intensity is independent of proliferation by showing that, whereas about 80% of L-plastin+ cells have “hyper long” telomeres in a young WT gut (**Fig 1E**), under 10% of these cells are proliferating, as assessed by double L-plastin/PCNA (Proliferating Cell Nuclear Antigen) staining (**Fig 1C**). Moreover, contrary to the differences between telomere intensities, there is no significant difference in proliferation between WT and *tert*^*-/-*^ immune cells (**Fig 1C1**).

**Fig 1.**
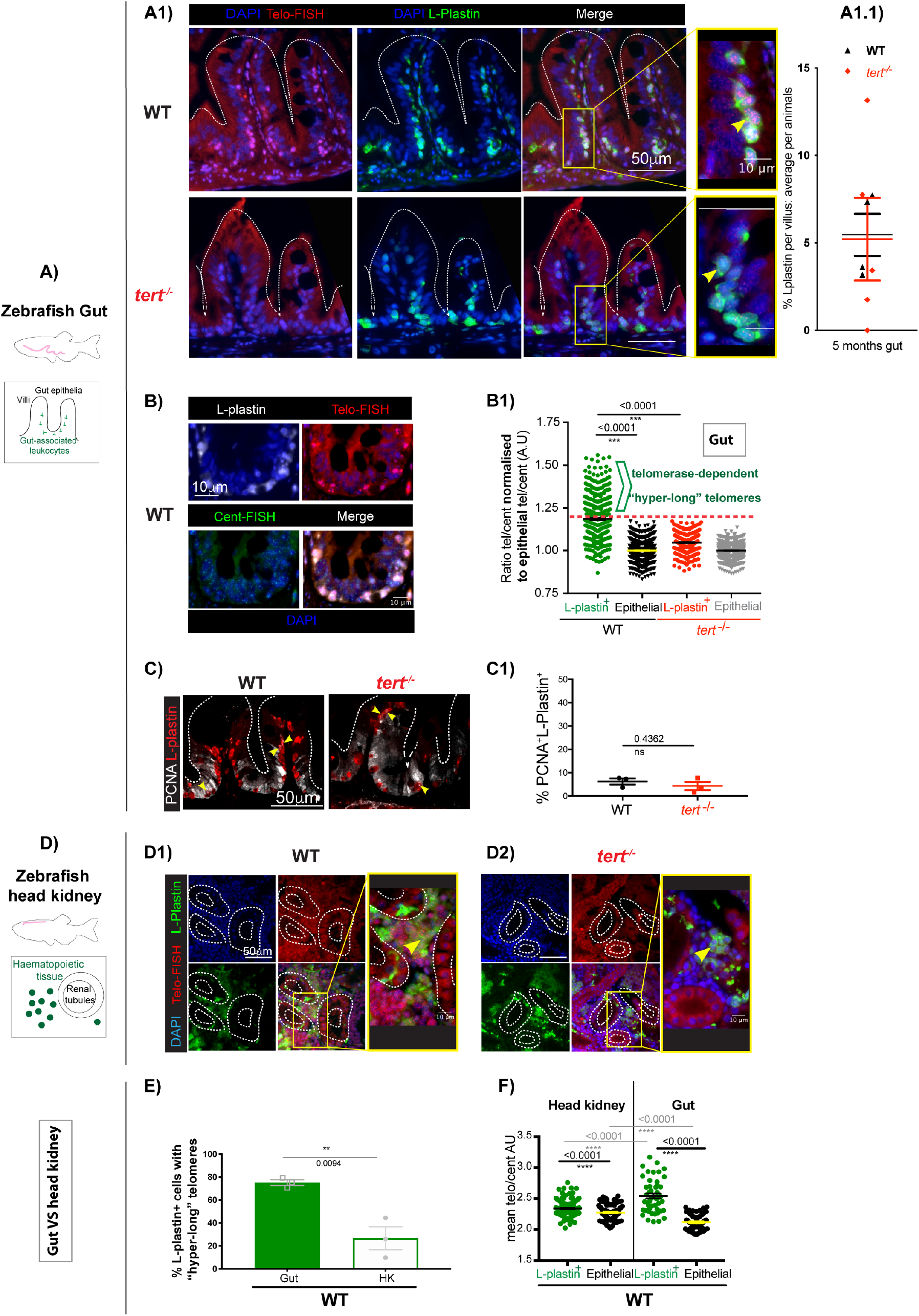
Gut-associated leukocytes have telomerase-dependent “hyper-long” telomeres, independently of proliferation: **A)** Zebrafish gut paraffin sections **A1)**, showing telomere TelC-Cy3 CCCTAA PNA probe in situ hybridisation (Telo-FISH) in red, combined with anti L-plastin immunofluorescence, in green. Nuclei counterstaining with DAPI in blue. Representative images shown from young (c.5 months) WT and *tert*^*-*/-^. **A1.1**) Quantifications of the average % of L-plastin+ cells per gut villi, average per animal. **B)** Representative images of Young WT gut with Telo-FISH combined with a near centromeric probe (ZEFRFAL1), which we call Cent-FISH. **B1)** Relative telomere length (Telo-FISH/Cent-FISH signals) quantifications defining the hyper-long telomere population as above 1.20 when normalised to the relative telomere length of nearby villi epithelial cells. **C)** Representative images of Young WT gut showing a double immunofluorescence staining against PCNA (white) and L-plastin (red) and **C1)** quantifications of the % of PCNA+L-plastin+. **D)** Zebrafish Head kidney paraffin sections, showing telomere TelC-Cy3 CCCTAA PNA probe in situ hybridisation (Telo-FISH) in red, combined with anti L-plastin immunofluorescence, in green. Nuclei counterstaining with DAPI in blue. Representative images shown from young (c.5 months) **D1)** WT and **D2)** *tert*^*-*/-^. **E)** Quantifications of the % of L-plastin^+^ with hyper-long telomeres comparing young WT gut with corresponding head kidney. **F)** Relative telomere length quantification in L-plastin and epithelial cells in gut and Head kidney. All scale bars: 50μm.

Even though the zebrafish gut share many similarities with the mammalian gut, including most of its diversity of immune sub-types^61,62^, there are still some key differences^62,63^. A relevant difference to have in mind for this study is that zebrafish do not have organised lymphoid structures, such as mesenteric lymph nodes (MLN), isolated lymphoid follicles (ILFs) or Peyer’s patches (PP). Moreover, the origin of resident gut macrophages, as well as the relative contribution of monocytes from the periphery towards the resident gut population is still largely unknown^61^. Whether there is a constant replenishment from the periphery or whether there is a localised immune stem cell pool or both, remains to be determined in zebrafish. Therefore we hypothesised that perhaps these “hyper-long” telomere immune cells had inherited these long telomeres because they had originated in the head kidney marrow. The head kidney marrow can be considered the equivalent of the human bone marrow, and it is the main source of immune cells in the adult zebrafish^64^. Importantly, it was previously shown that the zebrafish head kidney has predominantly long telomeres, whereas other tissues in zebrafish show a mixture of long and short telomeres^55,56^, supporting the hypothesis.

Our data show that immune cells (L-plastin^+^) also have significantly longer telomeres than epithelial cells in the head kidney (**Fig1D, E, F**). However, immune cells have significantly longer telomeres in the gut than in the head kidney, suggesting that, even if there is a contribution of immune cells from the periphery, there is further telomerase modulation in the gut (**Fig 1E, F**).

### 2.2 Most hyper-long telomere gut-associated leukocytes in zebrafish are macrophages

Since, akin to humans, the zebrafish gut contains cells from both innate and adaptive immune lineages^41,61,62,65-67^, we set out to identify the different subsets of immune cells from within the “hyper-long” telomere gut immune population (L-plastin^+^). To do this, we used a combination of available immune-specific reporter fluorescent transgenic zebrafish lines and immune-specific antibodies alongside Telo-FISH, in WT young adult tissue sections. In specific, we used what has been described as a macrophage-specific reporter (mpeg1.1:mcherry caax)^68^ and a neutrophil-specific reporter (*mpx*:gfp)^69^ transgenics. In addition, we used a zebrafish specific anti-T-cell receptor (TCR) antibody to identify T-cells (**Fig 2A**). These three lines allowed us to observe that while putative macrophages (*mpeg*+) and T-cells (TCR+) both have “hyper-long” telomeres, this is not the case for neutrophils (*mpx*+). To further validate the identity of these cells and be able to quantify the relative proportion of these immune subsets from amongst the “hyper-telomere” immune cell population, we used the double transgenic *mhcII*:gfp/*cd45*:dsred zebrafish line^70^. This line has been reported to allow the identification of macrophages/dendritic cells (*mhcIIdab*:gfp^+^*cd45*:dsred^+^); B-cells (*mhcIIdab:gfp*^*+*^*cd45:dsred*^-^) and T-cells/neutrophils (*mhcIIdab:gfp*^*-*^*cd45:dsred*^+^) in the zebrafish gut^70^ (**Fig 2B**). Using this transgenic line, we calculated that the relative proportion of immune subsets from within the “hyper-telomere” length population is constituted by c.60% putative intestinal mononuclear phagocytes (MPs) (Macrophages (MΦs/dendritics(DCs) (*mhcIIdab*:gfp^+^c*d45*:dsred^+^), c.10% B-cells (*mhcIIdab*:gfp^+^*cd45*:dsred^-^) and c.10% T-cells (*mhcIIdab*:gfp^-^*cd45*:dsred^+^) (**Fig2A and B**). Because some recent reports have highlighted that the *mpeg1*.*1* promoter in transgenic zebrafish may also mark some non-macrophage cells, such as a sub-population of B-cells in the gut^71^, we compared the result obtained from the *mhcII*:gfp/*cd45*:dsred transgenic line with the *mpeg1*.*1:mcherry caax* line and obtained similar results. In specific, we calculated that about 60% of “hyper-long” telomere cells were *mpeg*^+^ in the zebrafish gut (**Fig 2C**). Together, these two transgenic lines gave us the confidence to conclude that the majority the “hyper-long” telomere cells in the zebrafish gut are likely to be intestinal mononuclear phagocytes (MPs) or macrophages.

**Fig 2.**
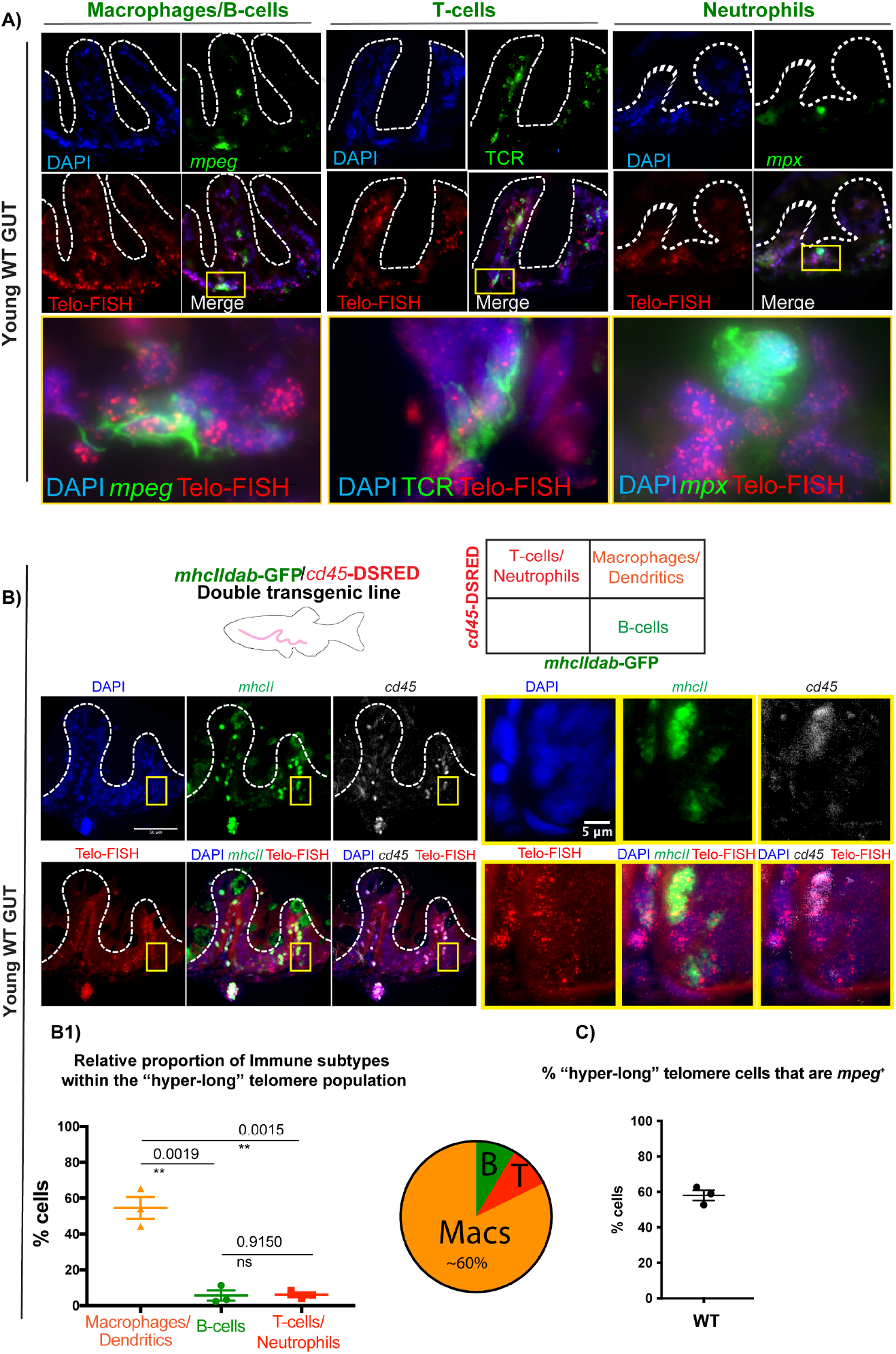
Most hyper-long telomere gut-associated leukocytes in zebrafish are macrophages. **A)** Telo-FISH (red) combined with anti-RFP antibody to detect *mpeg1*.*1:mcherry caax*^*+*^ cells (green), which have been described as mostly macrophages and a subset of B-cells; anti-TCR (green) to detect T-cells and anti-GFP to detect *mpx*:GFP+ expressing cells, which have been identified as neutrophils. **B)** *mhcIIdab*-GFP/*cd45*-DSRED transgenic line was used to perform Telo-FISH (red) combined with anti-GFP to detect *mhcIIdab*-GFP and anti-RFP to detect *cd45*-DSRED. Double *mhcIIdab*-GFP^+^/ *cd45*-DSRED+ have been identified as macrophages or dendritic cells; single *mhcIIdab*-GFP^+^ as B-cells and single *cd45*-DSRED as T-cells or neutrophils.**B1)** Quantification of the relative proportion of immune sub-types within the “hyper-long” telomere population, using the transgenic from B). **C)** Quantification of the % of “hyper-long” telomere cells that are *mpeg*^*+*^, using the mpeg1.1:mcherry caax transgenic from **A)**. All animals are WT for telomerase, and are of a young age (c.5 months). Nuclei are counterstained with DAPI (blue) and all scale bars represent 50μm.

### 2.3 A subset of WT gut *mpeg*^+^ cells express telomerase (*tert*) but the relative proportion of these cells decreases with ageing and is accompanied by telomere shortening

If indeed, as our data so far suggest, there is a further modulation of telomerase in gut immune cells, and that the majority of the “hyper-long” telomere cells in the gut are macrophages, then we would expect to detect telomerase expression in these cells. Additionally, since these “hyper-long” telomeres are telomerase-dependent, and telomeres have been shown to shorten with ageing in zebrafish^55^, then you would hypothesise that whatever telomerase expression may exist in these cells in a young WT gut, it is likely to decrease in ageing. To test these hypotheses, we used the *mpeg1*.*1:mcherry caax* line and combined anti-mcherry immunofluorescence with telomerase (*tert*) RNA fluorescent *in situ* hybridisation in young adult (c.5 months) and old WT (>36 months) zebrafish tissue sections. We also combined this with anti-PCNA immunofluorescence staining, to be able to distinguish between proliferating and non-proliferating cells, since you would expect that at least a subset of progenitor stem cells would express *tert*. (**Fig 3A**).

**Fig 3.**
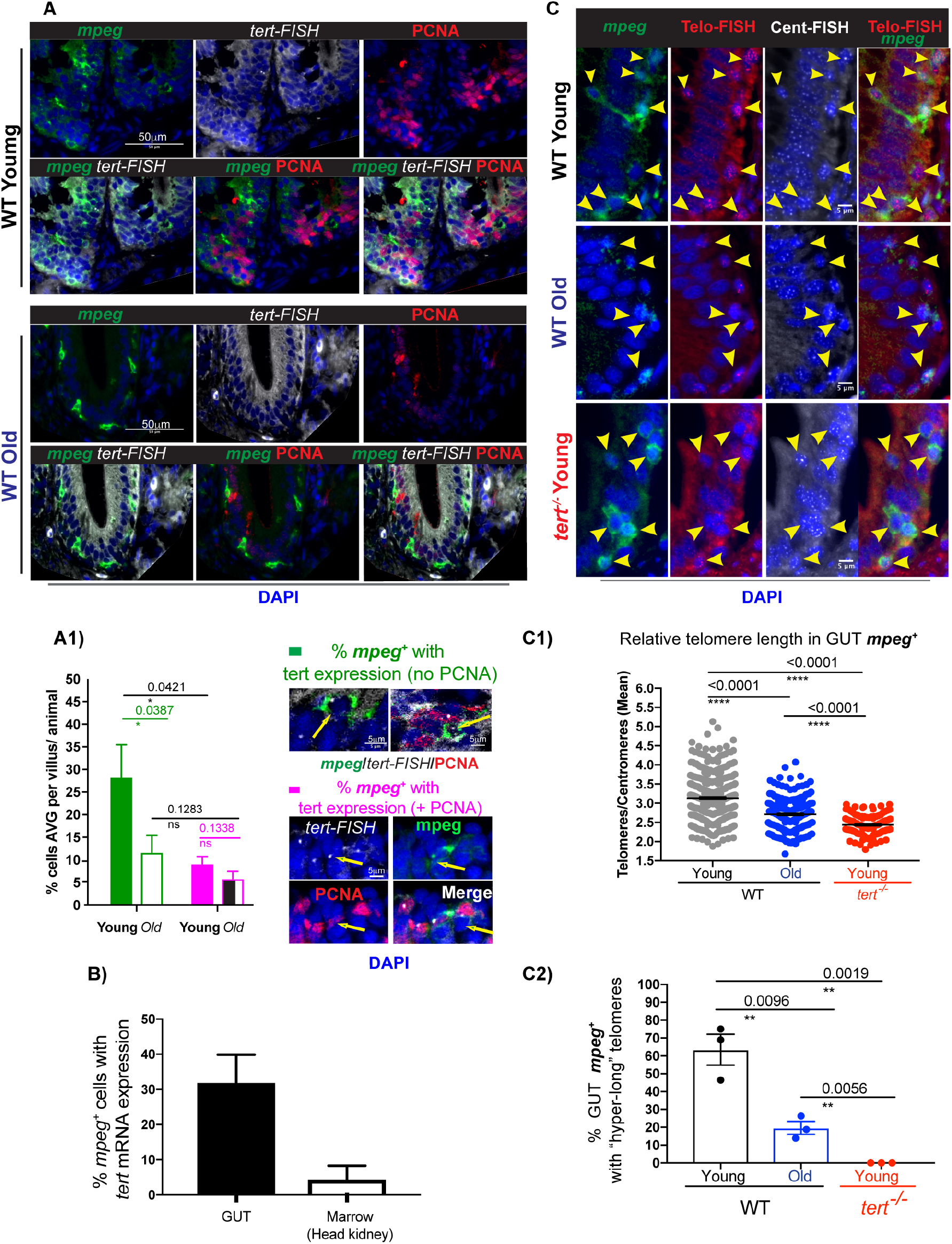
WT zebrafish gut *mpeg*^*+*^ cells express telomerase but this decreases with ageing and is accompanied by telomere shortening. The mpeg1.1:mcherry caax transgenic zebrafish line was used to **A)** detect telomerase (*tert*), by RNA *in situ* hybridisation (white) and PCNA expression in gut *mpeg*^+^ cells in young and old) WT zebrafish paraffin sections. **A1)** Quantification the % of *mpeg*^+^ with *tert* expression and no PCNA expression (green) and the % of *mpeg*^*+*^ cells with PCNA expression (magenta). In **B)** The relative % of *mpeg*^*+*^ cells with telomerase are compared between the gut and the head kidney of young WT fish. **C)** The *mpeg1*.*1:mcherry caax* transgenic zebrafish line was used to detect telomeres using Telo-FISH (red),Cent-FISH (white) combined with anti-RFP (in green) to detect *mpeg*^*+*^ cells young and old WT zebrafish paraffin sections. **C1)** Shows the relative quantification of telomere length (tel/cent ratio) in gut *mpeg*^*+*^ cells in WT young and old, compared to young *tert*^*-/-*^ zebrafish. **C2)** Shows the % of gut mpeg+ cells with “hyper-long” telomeres. Young animals are c.5 months old and old animals are >30-36 months old. Nuclei are counterstained with DAPI (blue). All scale bars represent 50μm.

Our data show that about 30% of *mpeg*^*+*^ cells express *tert* in WT young, compared to about 10% in WT old zebrafish gut (**Fig 3A1**), supporting the hypothesis that at least a subset of these cells expresses telomerase (*tert*) at a given point, and that the number of telomerase-expressing cells decreases with ageing. Accordingly, we observe a decrease in relative telomere length (telo/cent ratio) in gut *mpeg*^+^ cells with ageing (**Fig 3 C), C1)**) and a decrease in the proportion of *mpeg*^+^ cells with “hyper-long” telomeres (**Fig 3C2**). We also observe these decreases when analysing the bulk of gut immune cells as a whole (L-plastin^+^), rather than just *mpeg*^+^ cells (**Supp Fig 1**), suggesting that there is a general decrease in telomerase expression in gut immune cells with ageing (or a decrease in the proportion of immune cells that can up-regulate telomerase, with ageing). Relevantly, in young WT zebrafish gut, only about 10% of telomerase-expressing macrophages were proliferating (*mpeg*^*+*^ *tert*^*+*^ PCNA^+^) and we did not detect any significant difference in the proportion of these cells between young and old. Together, these data show that even though a proportion of gut *mpeg*^*+*^ cells express telomerase at young ages, the proportion of telomerase-expressing cells decreases with ageing, and is accompanied by telomere shortening. Decreased telomerase expression and telomere shortening in these cells seem to occur irrespectively of proliferation, suggesting that the expression of telomerase in these cells may have evolved to serve other non-canonical functions, rather than maintenance replicative potential.

### 2.4 Telomerase depletion accelerates age-associated increased apoptosis, decreased autophagy and decreased immune activation of zebrafish *mpeg*^*+*^ gut cells

Telomerase has been described to have both canonical (telomere elongation) and non-canonical (telomere elongation-independent) functions. We therefore set out to determine the key molecular changes occurring in gut *mpeg*^+^ cells in the absence of telomerase, at the young age of c.5 months. We chose molecular targets that would be expected to change in response to short telomeres, such as proliferation and DNA damage, as well as potential non-canonical changes, such as autophagy and immune activation markers. Importantly, we asked whether these alterations were also occurring with natural ageing and, if so, whether telomerase depletion accelerated such phenotypes, as this would indicate that such ageing phenotype are telomerase-dependent. Our results show that there is a significant decrease in the numbers of *mpeg*^*+*^ cells in the gut in old age, but that this is not significantly accelerated in the absence of telomerase, at least at the young age that our work is focusing on. (**Fig 4A), A1)**). In accordance with our previous observation that telomerase expression does not significantly affect gut immune cell proliferation (**Fig 1C), C1)**), we also do not detect significant changes in gut *mpeg*^+^ proliferation in either *tert*^*-/-*^ or natural ageing (**Fig 4A), A2)**). Even though we observe an increase in DNA damage response markers, namely *γ*H2AX, with ageing, this is not accelerated in the absence of telomerase, in the young *tert*^-/-^ (**Fig 4B), B1)**). We do, however, observe a significant telomerase-dependent increase in cleaved caspase 3^+^/ *mpeg*^+^ cells with ageing, suggesting that there is increased telomerase-dependent apoptosis in these cells (**Fig 4C), C1)**).

**Fig 4.**
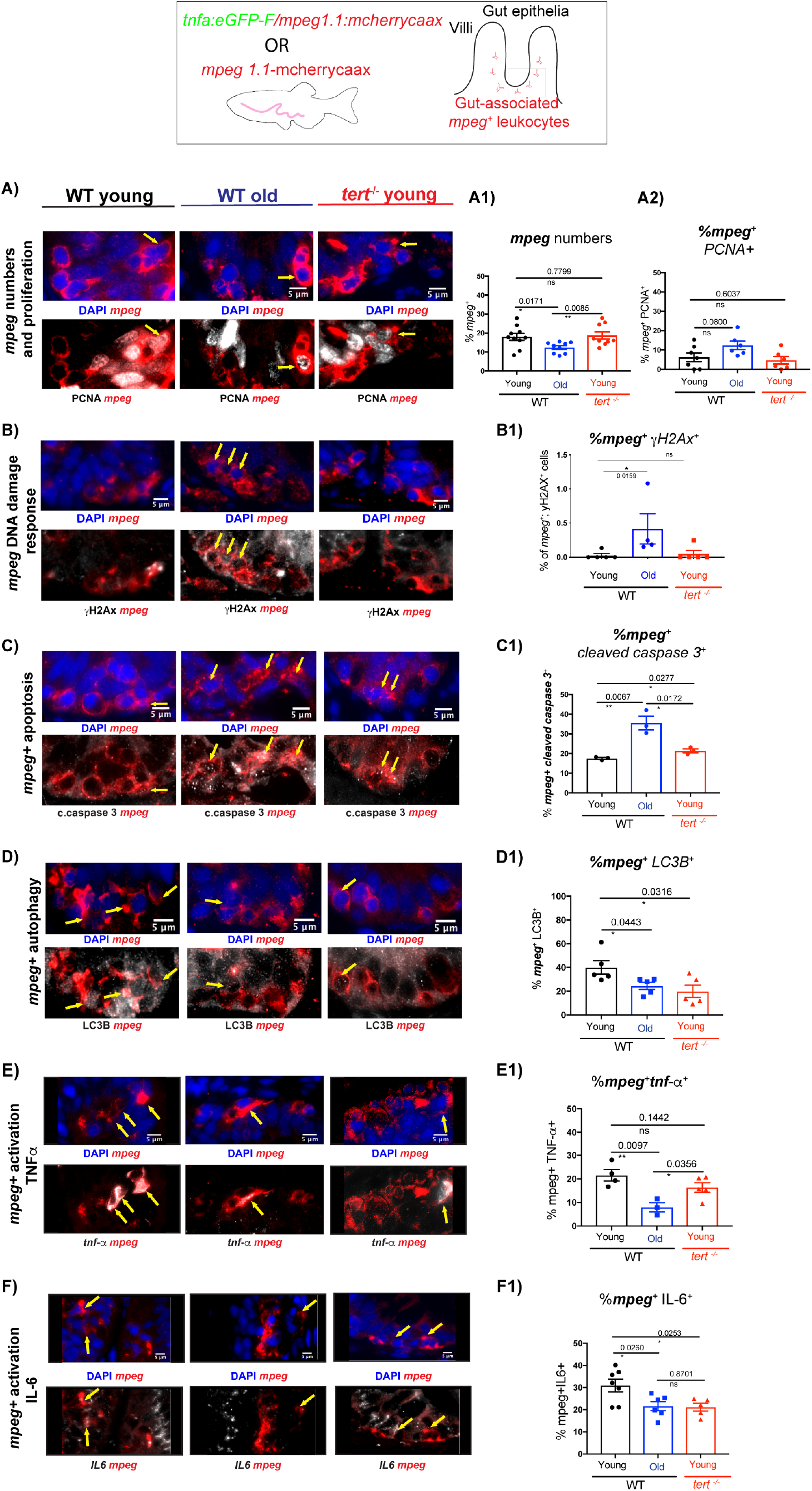
Telomerase depletion accelerates age-associated increased apoptosis, decreased autophagy and decreased immune activation of zebrafish gut *mpeg+* cells . The *mpeg1*.*1*:mcherry caax transgenic zebrafish line was used to assess **A)** numbers (**A1**) and proliferation (**A2**) of gut *mpeg*^*+*^ cells, using double immunofluorescence staining against RFP to detect *mpeg1*.*1:mcherry caax* (red) and anti-PCNA (white) to detect proliferating cells. Using the same fish (mpeg1.1:mcherry caax), we further analysed: **B**) DNA damage response (DDR), using double immunofluorescence against RFP to detect *mpeg1*.*1:mcherry caax* (red) and *γ*H2AX (white) to detect DDR. **B1)** Quantification of the % of *mpeg*^+^ *γ*H2AX^+^ cells; **C)** Apoptosis, using double immunofluorescence against RFP to detect *mpeg1*.*1*:mcherry caax (red) and anti-cleaved caspase 3 (white) to detect apoptotic cells. **C1)** Quantification of the % of *mpeg*^+^ cleaved-caspase 3^+^ cells; **D)** Autophagy, using double immunofluorescence against RFP to detect *mpeg1*.*1*:*mcherry* caax (red) and LC3B (white) to detect autophagosomes. **D1)** Quantification of the % of *mpeg*^+^ LC3B ^+^ cells. *tnf*a:eGFP-F/*mpeg1*.*1*:mcherry *caax* fish were used to assess immune activation, **E)** using double immunofluorescence against RFP to detect *mpeg1*.*1*:mcherry (red) and anti-GFP (white) to detect *tnfa* expressing cells. **E1)** Quantification of the % of *mpeg*^+^ *tnfa* ^+^ cells; **F)** using double immunofluorescence against RFP to detect *mpeg1*.*1*:mcherry *caax* (red) and anti-IL-6 (white) to detect IL-6 expressing cells. **F1)** Quantification of the % of *mpeg*^+^ IL-6 ^+^ cells. All animals are WT for telomerase, and are of a young age (c.5 months). Nuclei are counterstained with DAPI (blue) and all scale bars represent 50μm.

Because autophagy is a key molecular mechanism for immune function and telomerase has been shown activate autophagy in different cells types^72-75^, we hypothesised that gut macrophage autophagy would be affected in the absence of telomerase. Our data show that there is a significant decrease in autophagosomes, as assessed by LC3B puncta, in the *mpeg*^*+*^ gut cell population with old age, which again is accelerated in the absence of telomerase (**Fig 4D), D1)**). This suggests that telomerase contributes to the regulation of autophagy in gut *mpeg*^*+*^ cells and telomerase depletion is likely to contribute to autophagy defects in these cells in old age. Finally, we tested for the expression of key immune activation markers, known to be important for gut macrophage function, namely TNF*α* and IL-6. For this, we used the *mpeg*^*+*^*mcherry* line crossed with the *TgBAC(tnfα:GFP)*^48^ and an anti-IL-6 antibody. Our data show that old age is associated with decreased numbers of *tnfα* expressing *mpeg*^*+*^ cells, as well as a decreased numbers of IL-6 expressing *mpeg*^+^ cells (**Fig 4E), E1), F), F1)**). In specific, we further show that decreased numbers of IL-6 expressing *mpeg*^+^ cells is accelerated in the absence of telomerase (*tert*), suggesting that this is a telomerase-dependent mechanism.

### 2.5 Telomerase depletion leads to impaired phagocytosis in *mpeg*^+^ cells *in vivo* and increased gut permeability

As the name indicates, a key role, even if not the only one, for intestinal mononuclear phagocytes (MPs) is to phagocytose^40^. A remarkable characteristic of gut MPs is their ability to phagocytose foreign material without generating an inflammatory response, and gut macrophages do not fall within the M1/M2 classic phenotypes. This ability is essential in their role in discriminating between pathogens and other non-harmful antigens, such as food and microbiota^40^. It was therefore difficult to predict, whether the changes observed in the numbers of *mpeg*^+^*tnfα*^+^ and *mpeg*^+^IL-6^+^ with ageing, and, in the case of IL-6, accelerated in the *tert*^*-/-*^, would lead to changes *mpeg*^+^ function. Nevertheless, both IL-6 and TNF*α* have been described to play a role in macrophage’s immunesurveillance ability ^76,77^. To test whether immunesurveillance was affected in the zebrafish gut with old age, and whether this was telomerase-dependent, we adapted the well-described *E*.*coli* phagocytosis assay, using pHrodo™ Green *E. coli* BioParticles™ Conjugate for Phagocytosis^70^. We adapted this assay to assess phagocytosis specifically in the gut, *in vivo*. For this, we optimised delivery of these particles via oral gavage in adult *mpeg*^*+*^*-mcherry caax* zebrafish, both at young (c.5 months) and old (c. 35 months) WT ages, and in the absence of telomerase (*tert*^*-/-*^) at c.5 months. *mpeg*^*+*^ cells can be detected by the membrane bound *mcherry caax* and the % of *mpeg*^*+*^ cells containing visible green *E*.*coli* bioparticles inside (the zebrafish gastrointestinal tract is not acidic^62^, so the phRodo moiety ensures that these particles increase their fluorescence once phagocytosed into acidic vesicles) were quantified, as a readout for phagocytosis efficiency (**Fig 5A) A1)**). Our data show that whereas there is a trend towards decreased phagocytosis efficiency in WT old fish (p=0.0597) there is a clear statistically significant difference in phagocytosis efficiency in the absence of telomerase (p=0.0341) (**Fig 5 A2)**). This suggests that telomerase is required for efficient gut *mpeg*^+^ phagocytosis.

**Figure 5:**
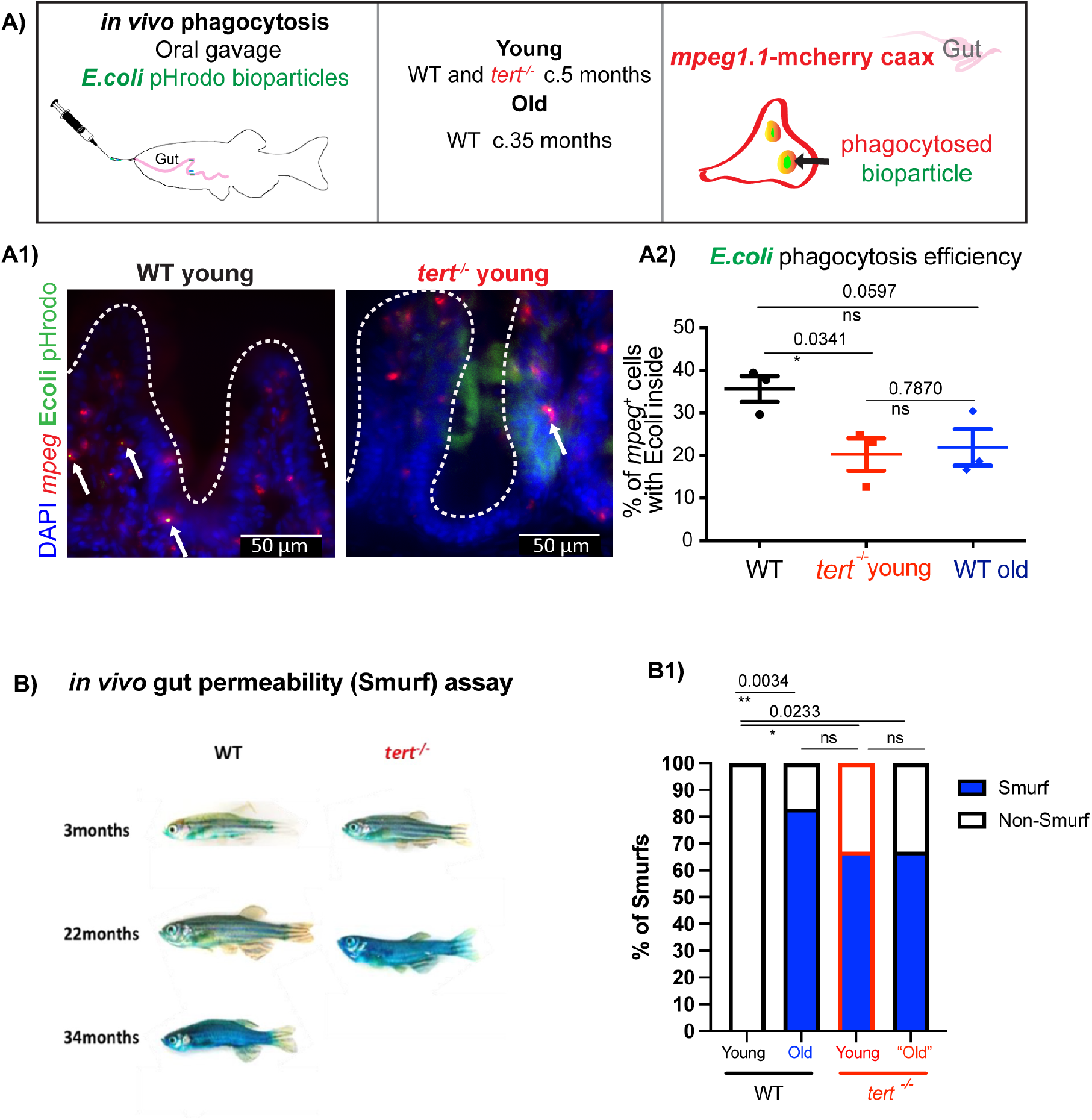
*In vivo* functional assays showing that telomerase depletion leads to impaired phagocytosis in gut *mpeg+* cells and increased gut permeability. The *mpeg1*.*1:mcherry caax* transgenic zebrafish line was used to quantify **A***) in vivo* phagocytosis in the gut, of *E*.*coli* pHrodo (green) bioparticles by *mpeg*+ cells (red). *E*.*coli* pHrodo bioparticles were delivered to animals’ stomachs via gavage. Phagocytosed E.coli pHrodo bioparticles will fluoresce green inside *mcherry caax*^*+*^ phagosomes, inside *mpeg*^+^ cells. **A1**) Shows representative images of young (c.5 months old) WT or *tert*^*-/-*^ zebrafish gut cryostats sections, displaying native *mcherry* (*mpeg*^*+*^ cells) and native GFP (*E*.*coli* pHrodo bioparticles). Nuclei were counterstained with DAPI (blue), and scale bars represent 50mm. **A2**) Shows quantification of the % of *mpeg*^+^ cells with *E*.*coli* bioparticles inside, as a readout of phagocytosis efficiency. B) *in vivo* gut permeability assay (Smurf assay). WT and *tert*^*-/-*^ of different ages were assessed for gut permeability by adding blue dye to the water for 30 minutes and then washing out extensively. Any fish developing an extensive blue coloration in their body indicate that the blue permeated through the gut, and were therefore considered “smurfs”. **B1**) Quantification of the percentage of “smurfs” (blue bars) per genotype. Young animals are c. 5 months old, old WT are >30 months old and “old” *tert-/-*are >12 months old, as further explained in materials and methods.

Cytokine expression in the gut is key to maintain gut homeostasis and regulate permeability, and most cytokine expression comes from innate immune cells. Because we observed a decrease in gut *mpeg*^+^ cells expressing *tnfα* and IL-6 with ageing, and decreased expression in IL-6, in particular, was accelerated in the absence of telomerase, we set out to test whether telomerase depletion affected gut permeability, a well described phenomena in old age and predictor of mortality in different organisms, including zebrafish^78^. Both IL-6 and TNFα expression by gut macrophages performs more complex functions than just being a pro-inflammatory cytokine. TNFα has been described to be involved in the regulation of enterocyte growth and can affect gut permeability^79^. TNFα can stimulate the production of matrix metalloproteinases and other tissue remodelling enzymes in the intestinal mesenchymal cells, central to regulating epithelial cell function^80^. Alternatively, IL-6 and TNFα expressing macrophages can contribute to the development of pro-inflammatory IL-17 expressing T-helper cells^81^. Previous work, including our own, had shown that gut permeability increases in naturally aged zebrafish. Whether telomerase played a role in this, remained to be determined. We used the well-described Smurf assay^78,82,83^, and show that indeed, depletion of telomerase accelerated the increase in gut permeability observed in old age (**Fig 5B), B1**), as assessed by the proportion of blue fish (Smurfs), in agreement with previous reports^78,83^.

## 3 DISCUSSION

The specific cellular and molecular mechanisms driving changes in immunity and inflammation in the gut with ageing, and their relative contribution to the clinical manifestations of an aged gastrointestinal tract are still poorly understood^47^. Telomere dysfunction is a primary hallmark of ageing, and there is evidence suggesting that the gut is one of the first tissues to age in a telomerase-dependent manner, with potential consequences for systemic ageing^54,55,84^. However, the consequences of ageing and, in specific, of telomere dysfunction in the immune component of the gut remain largely unexplored.

We set out to determine the molecular and functional consequences of whole organism telomerase depletion to gut-associated immunity, using the zebrafish model ^55-57^. Zebrafish has been established as an important complementary model of ageing, since, similarly to humans, requires telomerase for health and lifespan^55,56,58,85^. In this work, we show that gut-associated immune cells have telomerase-dependent “hyper-long” telomeres. In fact, the only significant difference in telomere length between WT and *tert*^*-/-*^ zebrafish at a young age (c. 5 months) is the absence of these telomerase-dependent “hyper-long” telomere cells (**Fig 1** and ^55,56^). We therefore postulated that these constituted a cellular subset particularly dependent on telomerase, and were therefore a potential candidate for driving the initial stages of telomerase-dependent ageing phenotypes, i.e, that are accelerated in the absence of telomerase, in the young *tert*^*-/-*^ model^55^. We therefore focused our study on the young *tert*^*-/-*^, and compared it with young and naturally aged WT, to address what molecular and functional changes occurring with age were already accelerated in the absence of *tert*^*-/-*^ at a young age, before the most severe gut degenerative phenotypes were described^55,56^.

Here, we show that gut “hyper-long telomere” immune cells are predominantly macrophages (*mpeg*^*+*^ or *mhcII*^*+*^/cd45+). Nevertheless, due to limitations in the availability of cell-surface immune-specific antibodies that work in zebrafish, we cannot exclude that we may be missing other specific subsets. In accordance with the presence of “hyper-long” telomeres, we show that a subset of gut macrophages (*mpeg*^+^) expresses telomerase in young WT zebrafish. Importantly, though, the relative proportion of telomerase-expressing macrophage*s* cells decreases with ageing and this is accompanied by an overall decrease in relative telomere length. This occurs in *mpeg*^+^ cells, but also in immune cells in general (L-plastin^+^). Together, this data suggest that whatever telomerase is expressed in gut immune cells and, in specific, *mpeg*^*+*^ macrophages, is not sufficient to prevent telomere shortening over time. This may be because the expression of telomerase decreases or, it may be that there is a progressive inability of activating telomerase with ageing, as has been described to be the case for memory T-cells, which become defective in their ability to proliferate with ageing^14,86^. However, in the case of gut mpeg^+^ cells, or indeed of gut immune cells in general, only a very small proportion of cells is proliferating at a given moment (c.10%). Moreover, this is not significantly affected with ageing or absence of telomerase. How are these cells shortening their telomeres then, and what do they need telomerase for? It may be that, even though only a small proportion of the immune cells are proliferating at any given time, this is still enough to contribute to telomere shortening. In fact, when we compare the telomere shortening in these cells with the telomere shortening in epithelial cells, which are constantly derived from proliferating progenitors, we observe that immune telomeres remain much longer than those in epithelial cells, even at older ages (**Supp Fig 2**). It may also be that decreased autophagy in gut immune cells, or at least, as our data suggest in *mpeg*^*+*^ cells, is leading to an accumulation of oxidative stress in these cells, which is a known contributor to rapid telomere attrition^87^. In fact, it has been shown that late generation telomerase null mice display altered mitochondria metabolism and increased oxidative stress in lung macrophages^88^. This could contribute to a much more significant telomere shortening than you would expect from a linear replicative telomere attrition, even in low proliferating cells^87^. It is also of note that, even though *tert*^*-/-*^ gut immune cells’ (L-plastin^+^) relative telomere length is much shorter, they are still statistically significantly longer than *tert*^*-/-*^ gut epithelial cells. This could potentially be due to alternative mechanisms of telomere length (ALT) being present, which have been suggested to co-exist alongside telomerase in zebrafish^89,90^. It could also simply reflect the fact that *tert*^*-/-*^ derive from a *tert*^+/-^ in-cross, since *tert*^-/-^ zebrafish are infertile^56,58^, therefore meaning that there is maternal contribution of telomerase expression during development. If these immune cells are seeded in the gut during development, as is the case for other tissues resident immune cells, such as the liver^91^, and given that they are lowly proliferating, they may have maintained relatively longer telomeres which may be sufficient to maintain relatively longer telomeres in low proliferating cells. Future work performing cell lineage tracing experiments will be required to determine the origins and maintenance mechanisms of adult, gut-associated immune cells.

Telomerase expression in innate immune cells has been shown to correlate with poor prognosis in age-associated, inflammation-driven diseases such as atherosclerosis^19^ and in unstable coronary plaques^20^. Furthermore, a study showed that transient transfection of macrophages with telomerase was sufficient to up-regulate matrix metalloproteinase (MMP) secretion, contributing to the tissue-damage characteristic of this disease^18^. Together, these data suggest that telomerase can contribute to the chronic activation and survival of macrophages thereby contributing to inflammation-driven diseases. In contrast, however, telomerase expression could also contribute to an anti-inflammatory state, by maintaining long-telomeres. When processed during DNA replication, telomeric sequences can bind to cytoplasmic Toll-Like Receptor 9 (TLR9), blocking downstream activation of pro-inflammatory pathways^92^. Indeed, recent work shows that synthetic telomeric sequences can modulate cellular energetics and can shift mice macrophages into a metabolically quiescent state, via mTor regulation^93^.

Our data show that, in the absence of telomerase, there is a variety of molecular mechanisms that are altered in gut *mpeg*^+^ cells. Some of these mechanisms are what you would expect from depletion of telomerase. In specific, we observe a decrease in telomere length over-time in gut *mpeg*^+^ cells increased DNA damage response and apoptosis with ageing. However, the increase in DNA damage response is not accelerated in the young *tert*^-/-^, suggesting that, at least at this age, and despite the shorter telomeres, it is still not sufficient to accumulate significant, unresolved, DNA damage. Moreover, we do not observe significant telomerase-dependent decrease in gut macrophage (*mpeg*^+^) proliferation with ageing, which would be another obvious consequences of telomerase-dependent telomere shortening. Some of the molecular changes we observe, however, are not necessarily the ones you would expect from telomere-shortening dependent, replicative senescence^5,6,85^. Instead, they may be better explained by non-canonical functions of telomerase. Telomerase is not only capable of restoring telomeres (canonical function) but also of performing non-canonical functions involved in the regulation of cell survival, inflammation and autophagy^72-75^ (recently reviewed here^21^). Here, we identified a telomerase-dependent decrease in autophagosomes (LC3B puncta) in gut *mpeg*^+^ cells with ageing. Additionally, we show that there is a decrease in *tnfα* and IL-6 expressing gut macrophages (*mpeg+)* with ageing, and that the decrease in *mpeg*^*+*^IL-6+ cells, in specific, is accelerated in the absence of telomerase. One could then hypothesis that these are likely consequences of restriction of the non-canonical functions of telomerase. Determining the relative contribution of canonical versus non-canonical telomerase functions in gut macrophage function in ageing will however, require future studies.

*In vivo*, a key feature of gut immunity is the ability to phagocytose pathogens. Our data show that the phenotypes we observed in the absence of telomerase accompany phagocytic defects in gut *mpeg*^*+*^ cells *in vivo*. The fact that phagocytic defects are not as significant in old age suggest that there may be only partial dependency on telomerase for this function in ageing or, as our data suggest, hat there are still some *mpeg*+ cells expressing telomerase in old age. Indeed our data show that there is a c.50% reduction in telomerase-expressing mpeg+ cells, from c.40% to 20%, and this could explain this observation. If and which the telomerase-dependent deregulated mechanisms here described is the actual cause of impaired phagocytosis remains to be determined. Nevertheless, changes in autophagy and immune activation are good candidates for affecting phagocytosis efficiency. As an example, deregulated LC3-dependent autophagy has been reported to affect macrophage phagocytosis and polarisation (recently reviewed here^94^). In specific, autophagy is important in the recycling of cellular components and Adenosine Tri-phosphate (ATP), thereby contributing to the energy requirements of macrophage activation ^95^. In fact, inhibiting autophagy with 3-methyladenine (3-MA) led to decreased levels of IL-6 and TNF-α in mice^94^. Furthermore, the fact that autophagy and phagocytosis share the requirement of several genes in macrophages contributes to the hypothesis that a mechanism, such as telomerase, that affects one may also likely affect the other^96^. The molecular mechanisms regulating autophagy and phagocytosis in adult zebrafish gut macrophages, however, remain largely unknown, adding an extra challenge to dissecting the specific pathways responsible for this phenotype in this study.

Finally, we aimed to determine how the age and/or telomerase-dependent changes we identified in gut immunity could impact on the tissue degeneration with ageing. Key telomerase-dependent ageing phenotypes previously described in zebrafish include gut villi atrophy and accumulation of senescent cells. Additionally, previous studies also show increased thickness of the lamina propria, alongside augmented periodic acid Schiff (PAS) staining, both indicative of de-regulated inflammation^54,55,97^. However, TNFα and IL-6 expression by gut macrophages perform more complex functions than just being a pro-inflammatory cytokines. For example, TNFα can stimulate the production of matrix metalloproteinases and other tissue remodelling enzymes in the intestinal mesenchymal cells, central to regulating epithelial cell function^80^. TNFα has also been described to be involved in the regulation of enterocyte growth and can affect gut permeability^79^. Gut permeability has been highlighted as a key phenotype of ageing that closely associates with mortality. Accordingly, our data show an increase in gut permeability with ageing, and this was accelerated in the absence of telomerase, from an early age. However, our data show a decrease in gut *mpeg*^+^ cells expressing TNFα, with ageing, but this was not accelerated in the absence of telomerase, at the early ages tested. This suggests that the telomerase-dependent increase in gut permeability observed is likely not due to TNFα alterations in *mpeg*^*+*^ cells. It could still, however, be caused by a systemic increase in TNFα in circulation, which was not addressed in this study, but has been previously described to increase with ageing, in humans^98,99^ and mice^100^. Intriguingly, It is known that IL-6 can increased gut permeability by modulating tight junction permeability^101^. Our data, however, show a decrease in IL-6 expressing *mpeg*^*+*^ cells. We were not however capable of quantifying how much IL-6 was being produced by *mpeg*^*+*^ cells overall, so it is still a possibility that overall levels of IL-6 are increased. Increased IL-6 levels as has in fact been reported to increase in the serum of aged mice^100^. A further hypothesis for how the telomerase-dependent changes in gut immunity here described to contribute to increased gut permeability is that decreased immune activation leads to, not only impaired *E*.*coli* phagocytosis but also impaired clearance of senescent cells. Previous studies have shown that macrophages are capable of clearing senescent cells in some contexts (recently reviewed here^102^) and that, with ageing, macrophages themselves can acquire senescent-like markers, including an M2-like phenotype^103^. Additionally, senescent-like macrophages have also been shown to induce senescence in other cells in a paracrine fashion, known as the “bystander effect”, thereby capable of amplifying the accumulation of senescent cells *in vivo*^*104,105*^. The phenotypes we observe here are in fact consistent with an increased senescent macrophage pool, with decreased numbers of M1-like (*tnfa*^+^ and/or IL-6^+^). Accumulation of senescence has been previously described to occur with ageing, in zebrafish, in a telomerase-dependent manner^55^. The senescence associated secretory phenotype has been described to include matrix metalloproteinases (MMPs), which are potential candidates for tissue destruction and could be involved in affecting gut permeability^106^. Additionally, impaired pathogen clearance may also lead to changes in the microbiome, which has also been implicated in disrupting intestinal permeability^107,108^.

## 4 CONCLUSIONS

Together, our data show that limiting levels of telomerase contribute to alterations in gut immunity, impacting on gut permeability and the clearance of pathogens. We therefore highlight telomerase (tert) as a key regulator of gut immune functions, likely to contribute to the genesis of gut degeneration and, potentially, systemic alterations, with ageing. Future work using cellular specific manipulation of telomerase will be required to test the requirement and sufficiency of telomerase depletion (canonical and non-canonical) in gut immunity to the general local and systemic phenotypes of old age.

## 5 MATERIALS AND METHODS

### 5.1 Zebrafish husbandry

Zebrafish were maintained at 27-28°C, in a 14:10 hour (h) light-dark cycle and fed twice a day. All experiments were performed in Home Office approved facilities at the Bateson Centre, at the University of Sheffield. All animal work was approved by local animal review boards, including the Local Ethical Review Committee at the University of Sheffield (performed according to the protocols of Project Licence 70/8681 and PP445509.

### 5.2 Zebrafish strains, ages and sex

Four strains of adult zebrafish (*Danio rerio*) were used for these studies: wildtype (WT; AB strain), *tert*^*-/-*^ (*tert*^*AB/hu3430*^), *Tg(mpeg1:mCherryCAAX)sh378* and *Tg(mpx:gfp)(*Source: University of Sheffield); *TgBAC(tnfα:GFP)pd1028*^*48*^ (Source: Bagnat lab, Duke University, USA), crossed with the above mpeg Tg line to produce a double transgenic line (Source: University of Sheffield). All of mixed sexes. Wild-type (WT; AB strain) have been obtained from the Zebrafish International Resource Center (ZIRC). The *telomerase* mutant line *tert*^*AB/hu3430*^ was generated by *N*-ethyl-nitrosourea mutagenesis (Utrecht University, Netherlands; Wienholds, 2004). It has a *T*→*A* point mutation in the *tert* gene and is available at the ZFIN repository, ZFIN ID: ZDB-GENO-100412–50, from ZIRC. The fish used in this study are direct descendants of the ones used previously^29,30^, by which point it had been subsequently outcrossed five times with WT AB for clearing of potential background mutations derived from the random ENU mutagenesis from which this line was originated. The *tert*^*hu3430/hu3430*^ homozygous mutant is referred to in the paper as *tert*^*-/-*^ and was obtained by in-crossing our *tert*^*AB/hu3430*^ strain. Genotyping was performed by PCR of the *tert* gene^29,30^. The telomerase null mutant (*tert*^*-/-*^) zebrafish, extensively characterised elsewhere^29-31,72^, displays no telomerase activity and has significantly shorter telomeres from birth, ageing and dying prematurely^29^. While *tert*^*-/-*^ fish have a lifespan of c.12-20 months, WT fish typically die between 36-42 months of age^29,30^. In order to study age-related phenotypes in zebrafish, in this study we use an age >30 months old fish for what we consider old in WT (in the last 25-30% of their lifespan), and we consider the *tert*^*-/-*^ old fish at the equivalent age (>12 months) that corresponds to the last 25-30% of their lifespan, approximately, as has been described before^55^. In specific, ‘Old’ was defined as the age at which the majority of the fish present age-associated phenotypes, such as cachexia, loss of body mass and curvature of the spine. These phenotypes develop close to the time of death and are observed at >30 months of age in WT and at >12 months in *tert*^*-/-*29,30^.

### 5.3 Tissue preparation: paraffin-embedded sections and cryosections

Adult fish were culled by overdose of MS-222, followed by immersion fixation in 4% paraformaldehyde (PFA) buffered at pH 7.0. Whole fish were then processed for paraffin-embedded sections or for cryosections. Importantly, all quantitative comparisons presented in figures were performed in paraffin sections. Cryosections were only used for TCR staining in Fig 3, since this antibody did not work in paraffin sections, in our hands.

#### 5.3.1 Paraffin-embedded sections

Whole fish were fixed in in 4% (PFA) buffered at pH 7.0, at 4°C for 48-72 h, decalcified 0.5M ethylenediaminetetraacetic acid (EDTA) at pH 8.0 for 48-72 h, and embedded in paraffin by the following series of washes: formalin I (Merck & Co, Kenilworth, NJ, USA) for 10 min, formalin II for 50 min, ethanol 50% for 1 h, ethanol 70% for 1 h, ethanol 95% for 1 h 30 min, ethanol 100% for 2 h, ethanol 100% for 2 h 30 min, 50:50 of ethanol 100% : xilol for 1 h 30 min, xylene I for 3 h, xylene II for 3 h, paraffin I for 3h and paraffin II for 4 h 30 min. Paraffin-embedded whole fish were then sliced in sagittal 3 μm-thick, using a microtome (Leica RM2265). These sections were then used for immunofluorescence, fluorescence RNA *in situ* hybridisation and Telo-FISH.

#### 5.3.2 Cryopreservation and cryosections

Dissected guts were washed in 1x phosphate-buffered saline (PBS), cut into anterior, distal and middle portions and were fixed in 4% PFA at 4°C, overnight (ON). Then, they were washed in cold PBS and immersed in 30% sucrose in PBS, ON at 4°C, for cryopreservation. Individual guts were then embedded in mounting media – optimal cutting temperature compound (OCT, VWR International), snap-frozen in dry ice, and stored at -20°C until cryosectioning. Cryosections were sliced at a 13 μm thickness using a Leica Jung Frigocut cryostat or a Leica CM1860 cryostat. Sections were air dried for 2 hours then used for immunohistochemistry or frozen for up to 3 months at -20°C before use.

### 5.4 Immunofluorescence (IF), telomerase (tert) RNA *in situ* hybridisation (RNA-ISH) and Telomere PNA FISH (Telo-FISH)

#### 5.4.1 IF

Before immunofluorescence staining, cryosections were hydrated in PBS at room temperature (RT) for 10 min, and paraffin-embedded sections were deparaffinised and hydrated as follows: histoclear (Scientific Laboratory Supplies, Wilford, Nottingham, UK) 2x for 5 min, followed by ethanol 100% 2x for 5 min, ethanol 90% for 5 min, ethanol 70% for 5 min, and distilled water 2x for 5 min. After antigen retrieval by sub-boiling simmering in 0.01 M citrate buffer at pH 6.0 for 10 min, using a microwave. After cooling, the sections were permeabilised in PBS 0.5% Triton X-100 for 10 min and blocked in 3% bovine serum albumin (BSA), 5% Goat Serum (or Donkey Serum), 0.3% Tween-20 in PBS, for 1 h. The slides were then incubated with the primary antibody at 4°C ON. After washes in PBS 0.1% Tween-20 (3x 10 min) to remove excess primary antibody, the sections were incubated with secondary antibody at RT for 1 h or at 4°C ON. Slides were then washed as above and incubated in 1 μg/ml of 4′,6-diamidino-2-phenylindole (DAPI, Thermo Fisher Scientific) at RT for 10 min. Finally, the slides were washed once in PBS, and mounted with vectashield (Vector Laboratories, Burlingame, CA, USA). The primary and secondary antibody details are described in **Tables 1 and 2**.

**Table 1.** Primary antibodies used for immunostaining.

| **Antibody,**<br>**species and type** | **Dilution**<br>**factor** | **Catalogue number;**<br>**Company, City, Country** |
| --- | --- | --- |
| PCNA rabbit polyclonal | 1:300 | GTX124496; GeneTex, Irvine, CA, USA |
| Rabbit anti-TCR-alpha (N terminus) | 1:200 | AS-55868, AnaSpec Inc |
| Mouse anti-RFP | 1:500 | GTX82561; Genetex |
| Chicken anti-GFP | 1:500 | AB13970, Abcam |
| Rabbit anti L-plastin | 1:300 | GTX124420, Genetex |
| Rabbit anti-LC3B | 1:200 | AB48394, Abcam |
| Rabbit anti-yH2ax | 1:300 | GTX127342, Genetex |
| Rabbit anti-IL-6 | 1:500 | AB6672, Abcam |

**Table 2.**
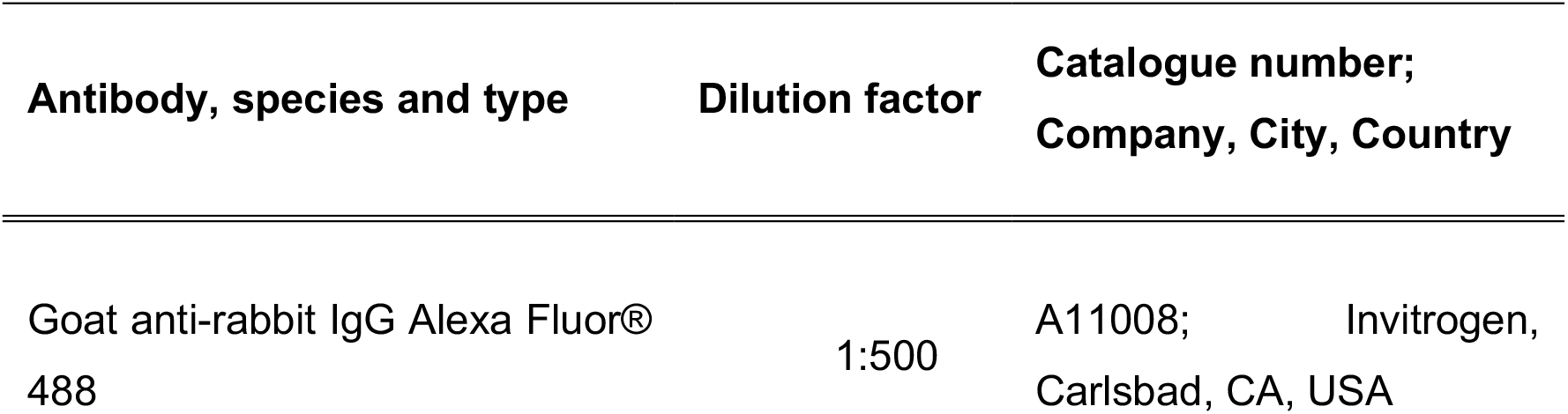
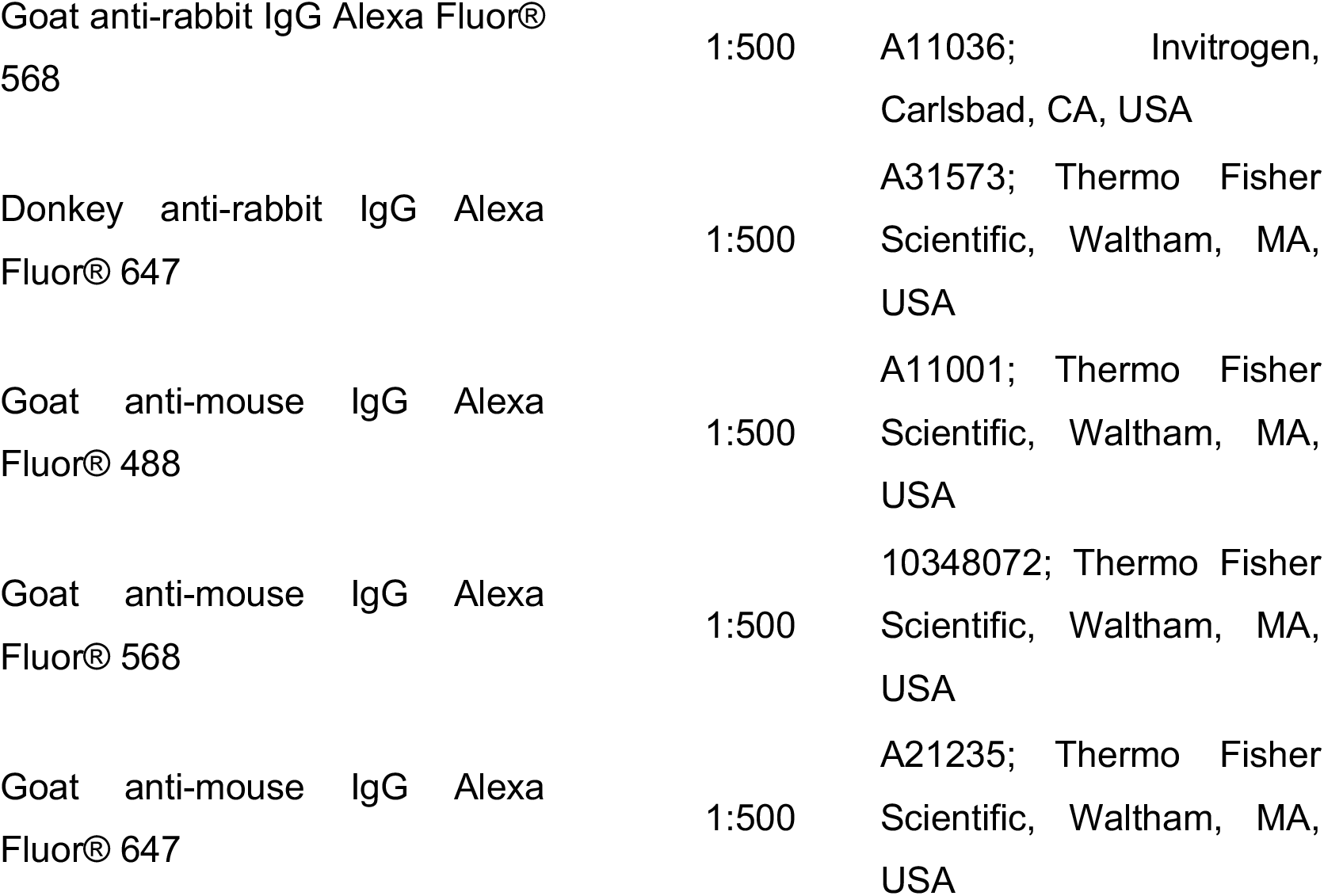
Secondary antibodies used for immunostaining.

#### 5.4.2 Telomerase (*tert*) RNA-ISH combined with IF

Paraffin sections were deparaffinised as described, using fresh solutions and bleach-treated glassware to avoid RNAse contamination. Where possible, solutions were made up in Diethyl pyrocarbonate (DEPC)-treated H_2_O or PBS and autoclaved. Following deparaffinisation, slides were rinsed in DEPC-PBS for 5 min and incubated at 37°C with 200ul of proteinase K at 20ug/ml in DEPC-PBS for 14 min (coverslipped). Slides were rinsed in DEPC-PBS briefly and fixed in 4% PFA for 20 min in a coplin jar. Slides were then rinsed in DEPC-PBS for 10 min at RT and acetylated as follows: 1ml DEPC-H_2_O, 11.2ul Triethanolamine (TEA) and 2.5ul acetic anhydride were made up per slide, immediately mixed and incubated on the slide for 10 min at RT. Slides were then rinsed in DEPC-PBS (10 min x2) and 1.5ml prewarmed hybridisation solution ((50% formamide, 2% Blocking Reagent, 0.1% Triton X-100, 0.5% CHAPS, 1mg/ml yeast RNA, 50ug/ml heparin (sodium salt) (all Sigma Aldrich), 5mM EDTA and 5xSSC pH7)) was incubated on the slides in a humidified box at 68°C for 1 hr minimum. Prewarmed digoxigenin-labelled probes (see below for probe information) were then added to slides (200ul per slide, at 2-5ng/ul in hybridisation solution) which were then coverslipped and incubated in a humidified box overnight at 68°C.Coverslips were removed and slides washed for 1hr at 68°C in a coplin jar in prewarmed ‘solution 1’: 25ml formamide, 12.5ml 20xSSC pH4.5, 5ml SDS, 7.5ml H_2_O, followed by 1hr at 68°C in prewarmed ‘solution 2’: 25ml formamide, 5ml 20xSSCpH4.5, 500ul Tween20, 19.5ml H_2_O. Gentle shaking was used during washes. Slides were rinsed twice with PBS+0.1% tritonX100 (PB-Triton), twice with maleic acid buffer (100mM maleic acid, 150mM NaCl, pH7.5) + 0.1% triton (MABT) then blocked in MABT+2% Blocking Reagent at RT for 1hr. Anti-DIG-peroxidase (POD) antibody was added at 1:500 in MABT+2% Blocking reagent, slides were coverslipped and incubated in a humidified box ON at 4°C. Slides were washed 6 × 20 min in PB-Triton in a coplin jar followed by 2 rinses in 100mM boric acid pH8.5+0/1% triton X-100. Fluorescent tyramides were incubated on the slides at 1:75 in freshly prepared TSA buffer (100mM Boric acid pH8.5, 0.1% Triton X-100, 2% dextran sulphate, 0.003% H_2_O_2_, 400ug/ml 4-iodophenol) for 20 mins at RT shielded from light. Finally, slides were rinsed 4x in PB-Triton, and fixed in 4% PFA for 5 min followed by a brief rinse in PBS. After RNA-ISH, immunofluorescence was performed on the same slides, avoiding exposure to light and starting from the permeabilisation step.

#### 5.4.3 *tert* probe design

Sense and anti-sense DIG-labelled RNA probes were synthesised from 5’ and 3’ regions of *tert* using DIG RNA Labelling Mix (Roche). A 562bp 3’ region of tert, was amplified using primers: tert 3’ FW, CGG TAT GAC GGC CTA TCA CT and tert 3’ REV, CAG GTT TTT TTT ACA CCC GC and TA cloned into PCRII (TA Cloning(TM) Kit Dual Promoter, Invitrogen). For antisense probe synthesis, the resulting construct was linearised with BamHI and transcribed with T7 RNA polymerase (New England Biolabs, (NEB)). For sense probe synthesis, the same plasmid was linearised with ApaI and transcribed with SP6 RNA polymerase (both NEB). ‘Full length’ tert cDNA was amplified using primers FW, ATGTCTGGACAGTACTCGAC and REV, CAGGTTTTTTTTACACCCGC. This was TA cloned as above into PCRII. A 1.5Kb subregion consisting of the 5’ end of this clone up to the first ApaI site was subcloned into PCRII and this 5’ tert plasmid was then used for probe synthesis of 5’ sense and antisense probes. For antisense, this was linearised with HindIII and transcribed with T7. For sense probe, it was linearised with ApaI and transcribed with SP6. A mixture of either 5’ and 3’ antisense or 5’ and 3’ sense (control) probes was used on sections in hybridisation buffer so that the final concentration of probe was 2-5ng/ul.

#### 5.4.4 IF combined with Telo-FISH

The IF protocol was followed as above. After the secondary antibody and PBS 0.1% Tween-20 washes, slides were washed three times in PBS and dehydrated with cold 70, 90 and 100% ethanol for 3 min each. Sections were denatured for 5 min at 80°C in hybridisation buffer (70% formamide (Sigma), 2.14mM MgCl2, 10mM Tris pH 7.2, 0.05% blocking reagent (Roche)) containing 0.5 ug/ml Cy-3-labelled telomere specific (CCCTAA) peptide nucleic acid probe (Panagene), followed by hybridization for 2 h at room temperature in the dark. The slides were washed once with 70% formamide in 2xSSC for 10 minutes, followed by 2෷10 minutes washes with 2x SSC. Sections were incubated with DAPI (SIGMA), mounted and imaged.

### 5.5 *In vivo* assays

#### 5.5.1 Phagocytosis

Delivery of pHrodo™ Green *E. coli* BioParticles™ (Thermo Fisher) to the gut was performed via oral gavage in adult *mpeg*^*+*^*-mcherry caax* zebrafish, both at young (c.5 months) and old (c. 35 months) WT ages, and in the absence of telomerase (*tert*^*-/-*^) at c.5 months. Zebrafish were not fed for 12/18hrs before 5 ul of E.coli bioparticles were delivered to zebrafish guts by oral gavage. Fish were sacrificed 4hours post gavage and gut tissue was dissected and processed for cryopreservation and sectioning. Cryosections of zebrafish gut were then imaged for native fluorescence, combined with DAPI nuclear staining. *mpeg*^*+*^ cells can be detected by the membrane bound *mcherry caax* and the % of *mpeg*^*+*^ cells containing visible green E.coli bioparticles inside (the zebrafish gastrointestinal tract is not acidic^62^, so the phRodo moiety ensures that these particles increase their fluorescence once phagocytosed into acidic vesicles) were quantified, as a readout for phagocytosis efficiency. We further ensured that the bioparticles were indeed inside the cells by going through all the Z-stacks, of 0.5mm each.

#### 5.5.2 Gut permeability

Smurf gut permeability was performed as described previously^83^, by placing zebrafish (WT and *tert*^*-/-*^) of different ages in individual tanks containing 2.5% (w/v) blue #1^78,83^ in water for 30 minutes. Individuals were then rinsed under clear water until no more blue colouration could be found in the eluate. Fishes showing extended coloration of blue in their body were considered as Smurfs.

### 5.6 Imaging and quantifications

Paraffin-embedded and cryosections sections were imaged by epifluorescence microscopy, using a DeltaVision microscope with a 40x oil objective. In order to quantify the alteration in the staining patterns, a z-projection was generated using ImageJ (Rasband, W.S., ImageJ, U. S. National Institutes of Health, Bethesda, Maryland, USA, https://imagej.nih.gov/ij/, 1997-2018.) at least 3 Fields of View (FoV) were imaged, containing at least two gut villi, per animal. At least three individual animals per genotype were imaged. Raw images were used for quantification and the images were then processed with Adobe Illustrator 21.0.2 for display purposes.

### 5.7 Statistical analysis

Statistics were performed using the GraphPad Prism v7.00. Normality was assessed by the Shapiro-Wilk test. For normally distributed data where most of the groups had a sample size of ≥5 unpaired t-test was used to compare 2 data points. For non-normally distributed data and/or data containing less than 5 animals in most of the groups Mann-Whitney test and Kruskal-Wallis tests were used instead. Chi-square was performed on the comparison between the number of Smurfs versus non Smurfs in the intestinal permeability assay. A critical value for significance of p<0.05 was used throughout all analysis. There were no repeated measurements performed in this study. Quantifications were either performed blind or/and by different individuals to increase robustness and confidence in the results obtained.

## SUPPLEMENTARY FIGURES

**Supplementary Figure 1:**
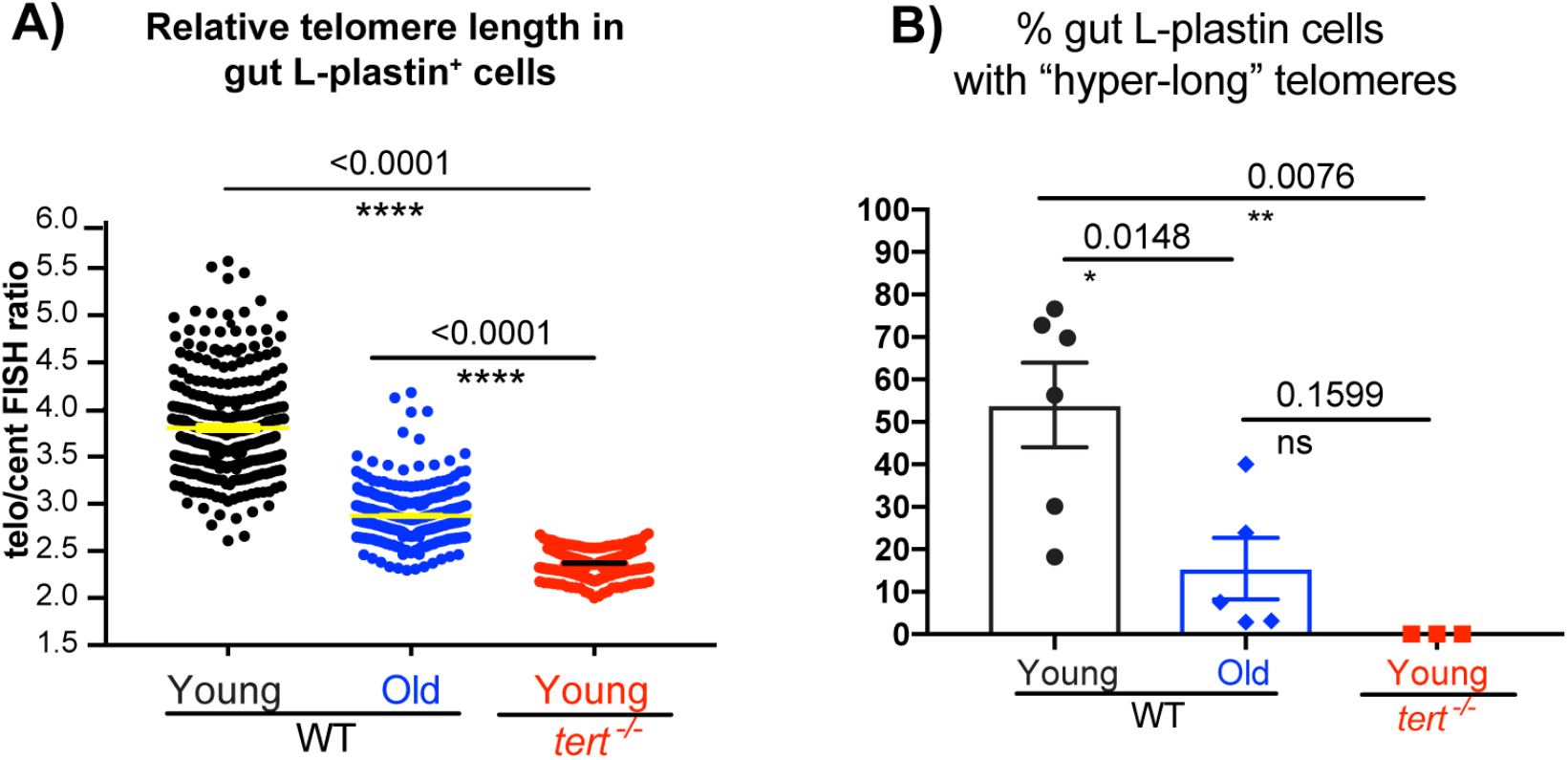
Telomere length decreases in gut immune cells with ageing. **A)** Quantification of the relative telomere length of L-plastin+ cells, from gut paraffin sections, using combine immunostaining for anti-L-plastin and telomere *in situ* hybridization (Telo-FISH), combined with the near-centromeric probe (Cent-FISH), as in Figure 1). **B)** From the same quantifications as in A), we can calculate the % of gut L-plastin+ cells with “hyper-long” telomeres. Young animals are c.5 months old and old animals are >30-36 months old.

**Supplementary Figure 2:**
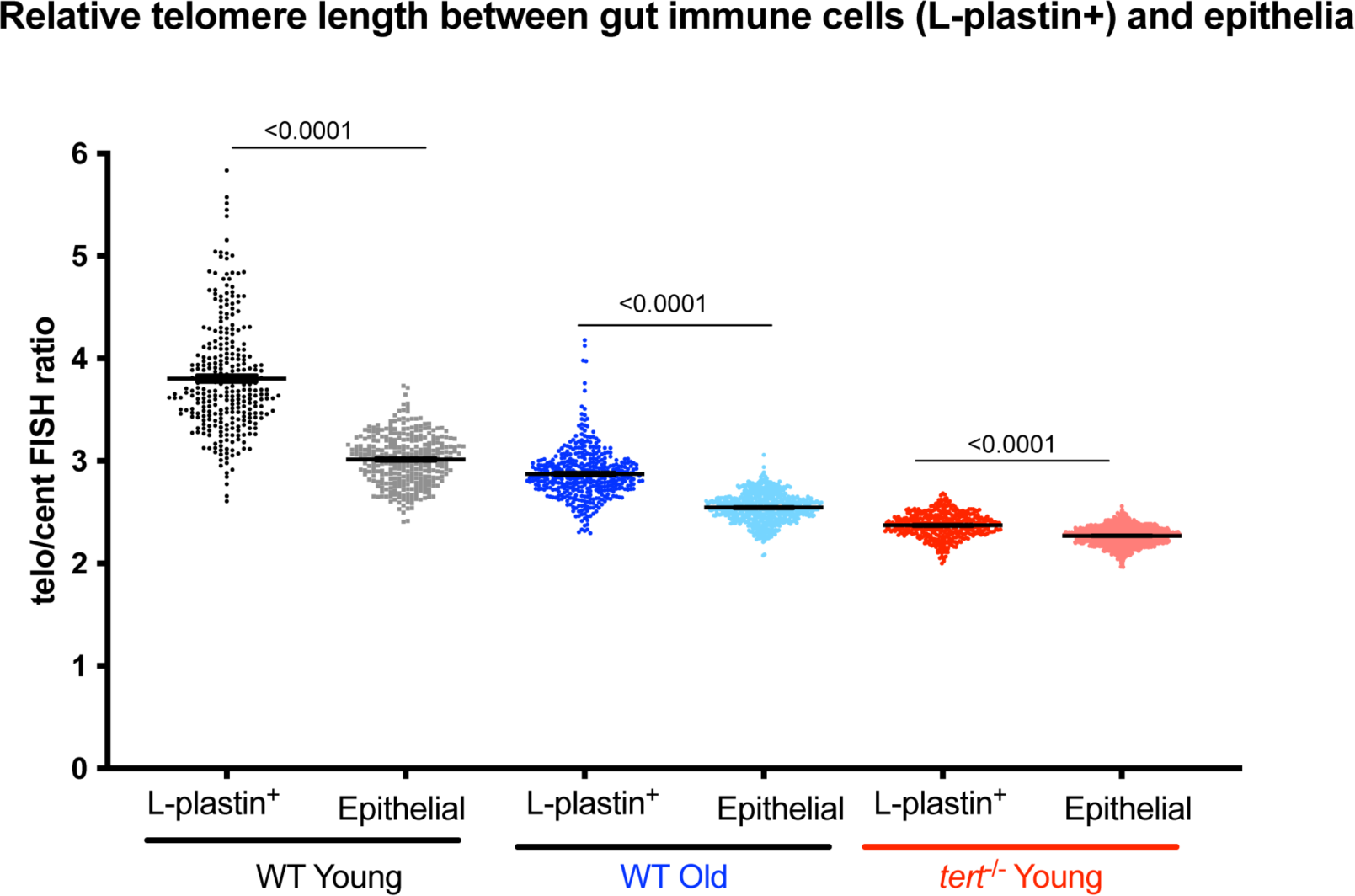
Immune cells in the gut retain longer telomeres than epithelial cells, despite telomere shortening over time. **A)** Quantification of the relative telomere length of L-plastin+ cells, from gut paraffin sections, using combine immunostaining for anti-L-plastin and telomere *in situ* hybridization (Telo-FISH), combined with the near-centromeric probe (Cent-FISH), as in Figure 1. Young animals are c.5 months old and old animals are >30-36 months old.

## ACKNOWLEDGEMENTS

We thank Dr. Valery Wittamer for kindly sharing the *mhcII*:gfp/*cd45*:dsred zebrafish transgenic line and Dr. Simon Johnston for sharing the TNFα:GFP crossed with the *mpeg1*.*1*:*mcherry caax* line, so that we could cross it back into the *tert*^-/-^ line. We also thank Dr. Simon Johnston and Dr. Daniel Humphreys for critical reading of the manuscript. We thank Prof. Ilaria Bellantuono for useful discussions during development of this work. Finally, we thank Dr. Miguel Godinho Ferreira for support during CMH’s post-doc, in which the work we did and publish together^55,56^ provided key insights that precluded the development of this work. We also thank Dr. Miguel Godinho Ferreira for sharing the *tert*^*-/-*^ line from the same outcross that had been used in the previous aforementioned studies.

## AUTHOR CONTRIBUTIONS

PSE helped design, conceived and analysed experiments; RRM helped design, conceived and analysed experiments; EJT performed and analysed experiments; AF performed and analysed experiments, SAR helped to conceive experiments, interpret results and provided the *mpeg1*.*1*-mcherry *caax* and the *mpx:GFP* transgenic lines; CMH conceived the study, designed, performed and analysed experiments, and wrote the manuscript with input from co-authors.

## DATA AVAILABILITY STATEMENT

The datasets and source data generated during and/or analysed during the current study are available from the corresponding author on reasonable request.

